# An evolutionary cell biology perspective into the diverging mechanisms of clathrin-mediated endocytosis in dikarya fungi

**DOI:** 10.1101/2024.03.28.587219

**Authors:** Andrea Picco, Christopher P. Toret, Anne-Sophie Rivier-Cordey, Marko Kaksonen

## Abstract

Clathrin-mediated endocytosis is an ancient eukaryotic trafficking pathway, which transports plasma membrane and associated cargo into the cell and is involved in numerous cell- and tissue-level processes. Cargo selection and clathrin-coated vesicle formation is mediated by over 60 proteins that assemble in a regular and sequential manner at the plasma membrane. Decades of endocytosis studies have followed the tenet that uncovering the conserved core molecular mechanisms is sufficient to understand a cellular process. However, this approach also revealed a number of cell type or species-related variations that challenge a universal conserved, core mechanism. In this paper, we refocus on the endocytic diversity to understand how evolution shapes endocytic mechanisms. We define a comparative evolutionary cell biology approach that uses dikarya fungi as a model clade and live-cell fluorescence microscopy to study endocytosis dynamics in three species: *Saccharomyces cerevisiae*, *Schizosaccharomyces pombe* and *Ustilago maydis*. Our results quantitatively define several phenotypic differences between the species. We uncover several differences that impact the endocytic early phase, the protein assembly order, actin regulation, membrane invagination and scission. These findings demonstrate a mosaic evolution of endocytic traits, suggest ancestral states and direction of changes. We also investigate the phenotypic plasticity and robustness against environmental conditions. Lastly, we demonstrate that relatively minor evolutionary changes can majorly impact endocytic phenotypes. These studies force an appreciation of endocytic variation as not auxiliary, but vital to mechanistic understanding of this conserved cellular pathway.

## INTRODUCTION

Cellular processes such as membrane trafficking, polarity establishment, cell division or cell migration take place across the diversity of eukaryotic life and are typically mediated by conserved protein machineries. Most contemporary cell biological studies emphasize the conserved aspects of these processes, and deemphasize the variation between species. In the conceptual framework of cell biology, the proteins and processes that are not broadly conserved are often considered as auxiliary for understanding the conserved “core mechanisms” of cellular processes. Eukaryotes, however, exhibit a wide range of diversity across their cellular processes. The molecular mechanisms behind this diversity and the evolutionary processes and pathways that lead to it remain poorly understood. The traditional conservation-focused perspective of cell biology needs to be complemented with an evolutionary perspective to reach a comprehensive understanding of the mechanisms, functions, and origins of cellular processes.

Clathrin-mediated endocytosis, is an ancient vesicular trafficking process that appeared with the emergence of eukaryotic cells [1] and occurs throughout the eukaryotic tree of life (ciliates, plants, amoeba, fungi, animals and many others [2–6]). The core function of endocytosis is to form membranous cargo-loaded transport vesicles from the plasma membrane [7]. Cargo sorting, regulatory mechanisms, and subsequent trafficking signals are coupled to endocytosis to maintain plasma membrane homeostasis, nutrient uptake and signaling in cells, which are contingent on species [8]. Furthermore, a number of physical parameters vary across species, and likely impact core mechanisms and machinery across species. Environmental and physiological parameters such as temperature, osmotic pressure, and membrane composition, also influence clathrin-mediated endocytosis [9,10]. To address the relationship between physiology, environment and evolution in the overall process requires comparable, well-defined model species.

The mechanisms of clathrin-mediated endocytosis have been primarily studied in mammals and in the two fungal species budding yeast, *Saccharomyces cerevisiae*, and fission yeast, *Schizosaccharomyces pombe*, [7,8,11]. Most endocytic proteins are conserved between mammals and the yeasts, and the sequential assembly of these proteins at endocytic sites is similar from the initial clathrin and adaptor recruitment to the final vesicle budding that coincides with an Arp2/3-mediated burst of actin polymerization [7]. Despite the similarities, there are also clear examples of endocytic mechanisms varying across eukarya. In mammalian cells, clathrin plays a central mechanistic role in membrane invagination, while actin polymerization can provide additional force for the final vesicle budding [7,8,12,13]. In contrast, actin provides the primary driving force for membrane invagination in fungi, while clathrin is dispensable for the process [14–16]. Moreover, there is evidence that even within fungi the mechanisms of endocytosis have diverged. A pioneering comparative study showed that protein dynamics and stoichiometries vary between *S. cerevisiae* and *S. pombe* [17]. These mechanistic differences underscore the importance of understanding, not only the conserved aspects, but also the variability of clathrin-mediated endocytosis.

It is challenging to decipher the evolutionary changes between distant lineages such as yeast and mammals, which have diverged for over a billion years and exhibit large physiological differences. Here, we define an evolutionary cell biology approach that uses three fungal species to study clathrin-mediated endocytosis. We use quantitative live-cell imaging to compare the dynamics of orthologous endocytic proteins in these three species.

## RESULTS

### An evolutionary cell biology approach for clathrin-mediated endocytosis

We chose three fungal species for our study: *S. cerevisiae, S. pombe,* and *U. maydis* (Figure 1A). These species are members of dikarya, a subkingdom of fungi that includes over 90% of known fungal species, and is subdivided into two main phyla. *S. cerevisiae* and *S. pombe* represent the ascomycota phylum and *U. maydis* the basidiomycota phylum. Ascomycetes and basidiomycetes diverged about 700 million years ago, which is comparable to the divergence time between *Drosophila melanogaster* and *Homo sapiens* (Figure 1A) [18,19]. These three fungal species share the ability to grow as single-celled yeasts (Figure 1A). They can be cultured in identical conditions, and are well-established genetic model organisms. Studies done on the two ascomycete species, *S. cerevisiae* and *S. pombe,* have revealed that endocytic proteins localize to the endocytic site with precise temporal dynamics, and the functional relationship between protein assembly and endocytic progression is well established [20–23]. In *S. cerevisiae,* over 60 endocytic proteins have been grouped into four functional modules: the endocytic coat, WASP/Myo (Arp2/3 regulators), scission proteins, and the actin filament network (Figure 1B) [20]. We chose to investigate eight endocytic proteins as representatives of the different endocytic protein modules defined in *S. cerevisiae*. We then used BLAST analysis to identify the orthologs in the other species (Figure 1C).

**Figure 1:**
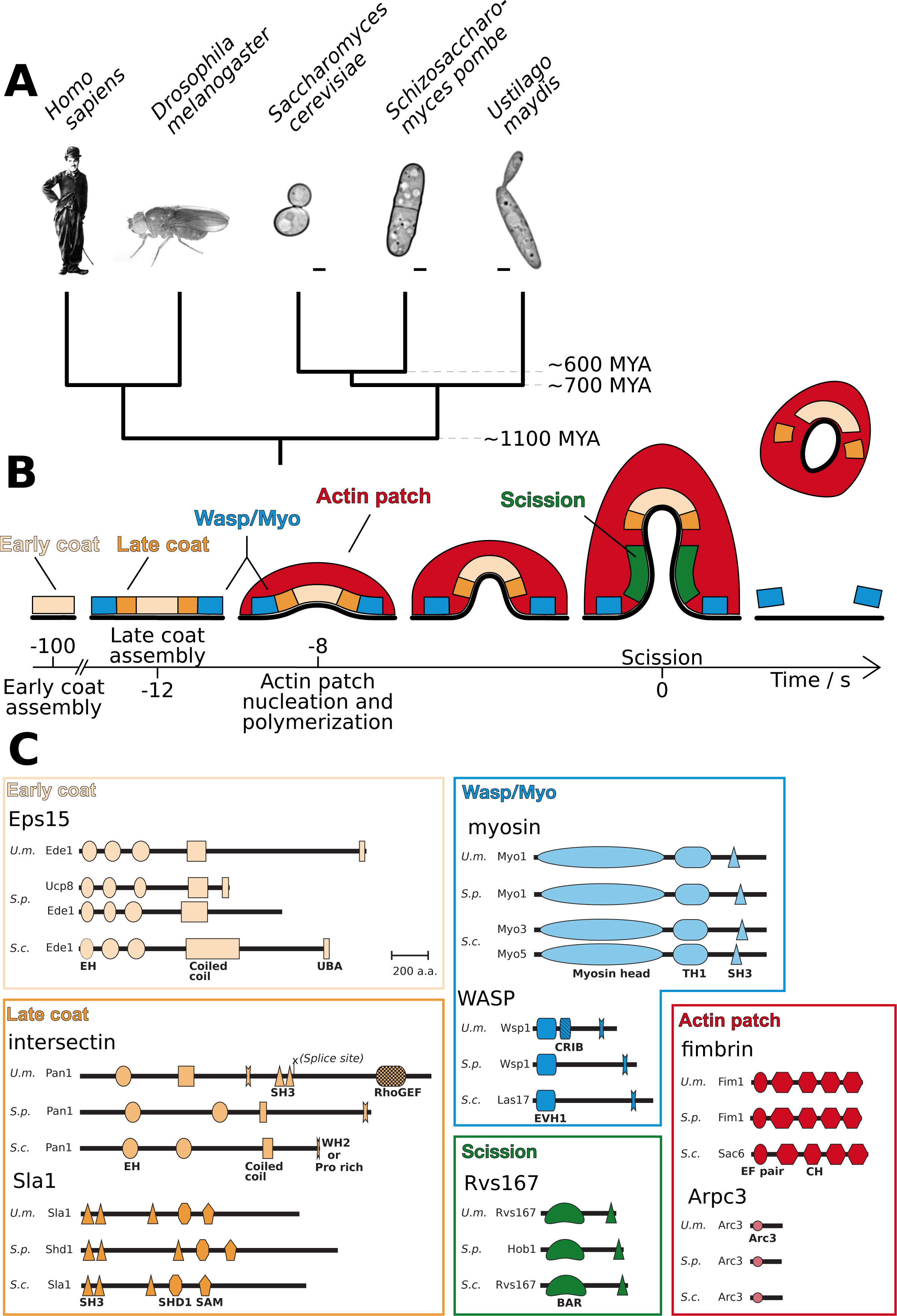
(A) A phylogenetic tree showing approximate divergence times between species. Yeast species are pictured with bright field images and are represented in scale. Scale bar = 1 μm. (B) Schematic depiction of the modular assembly of the *S. cerevisiae* endocytic pathway with approximate sequential timing and positions of each module relative to the plasma membrane (thick black line). Modules are color-coded. (C) Domain organization of different endocytic protein orthologs in the three species. Colors correspond to the endocytic modules represented in B.

The naming of *S. cerevisiae*, *S. pombe* and *U. maydis* orthologs are often inconsistent, or only systematic gene names exist. For simplicity, we refer to the proteins by the established names of their mammalian orthologs. The *S. cerevisiae* names are used for fungal-specific Sla1 and Rvs167 orthologs (Table 1).

**Table 1:** Endocytic protein orthologs.

| **Protein family** | **Mammalian orthologs** | ***Ustilago maydis***<br>**Systematic name**<br>**(Named in this study)** | ***Schizosaccharomyces pombe***<br>**Systematic name** | ***Saccharomyces cerevisiae***<br>**Systematic name** |
| --- | --- | --- | --- | --- |
| Eps15 | EPS15<br>EPS15R | UMAG_05533<br>(Ede1) | Ede1<br>Ucp8 | Ede1 |
| Intersectin | ITSN1<br>ITSN2 | UMAG_06013<br>(Pan1) | Pan1 | Pan1 |
| Sla1 |  | UMAG_05337<br>(Sla1) | Shd1 | Sla1 |
| WASP | N-WASP<br>WASP | UMAG_03687<br>(Wsp1) | Wsp1 | Las17 |
| Myosin I | MYO1E | UMAG_11115<br>(Myo1) | Myo1 | Myo3<br>Myo5 |
| Fimbrin | PLS1<br>PLS2<br>PLS3 | UMAG_04768<br>(Fim1) | Fim1 | Sac6 |
| Arc3 | Arpc3 | UMAG_10246<br>(Arc3) | Arc3 | Arc18 |
| Rvs167 | BIN1<br>BIN2<br>BIN3<br>AMPH | UMAG_01748<br>(Rvs167) | Hob1 | Rvs167 |

Ortholog names used in the manuscript.

From the early-arriving coat proteins we chose the Eps15 family orthologs, known as Ede1 in fungi. These orthologs are EH-domain-containing and ubiquitin-binding proteins in both *S. cerevisiae* and mammals, which promote endocytic site initiation [22,24,25] (Figure 1B and C). *S. pombe* has two Eps15 paralogs, suggesting that a gene duplication has occurred in the *S. pombe* lineage. One of the *S. pombe* Eps15 paralogs has lost the ubiquitin binding UBA domain, which suggests a possible separation of Eps15 functions (Figure 1C). From the late coat proteins, we chose the intersectin family proteins, known as Pan1 in fungi, and the fungal-specific Sla1 family proteins, which are late-arriving endocytic coat proteins with scaffolding and actin regulatory functions [26,27]. These late coat proteins move inward with the invaginating membrane at the end of the endocytic process [21,28] (Figure 1B). The ascomycete intersectins maintained only the N-terminal EH-domains and coiled-coil regions, while the *U.maydis* protein retains the SH3 and RhoGEF domains typical of the mammalian intersectin, but lacks the first EH domain (Figure 1C). We chose WASP and type I myosins to mark the WASP/Myo module. WASP is the predominant Arp2/3 activator of actin nucleation at endocytic sites [29,30]. One WASP ortholog is present in each species. *U.maydis* WASP contains a central regulatory CRIB domain, also present in animal WASP proteins, but lost in the ascomycetes (Figures 1B and C). The Class I myosins, which power internalization [31], are present in each species, but *S. cerevisiae* has a duplication (Myo3/5) (Figure 1C). To mark the endocytic actin network, we chose the fimbrins, which crosslink actin filaments [32], and Arc3, a subunit of the Arp2/3 actin nucleator complex [33]. The branched actin growth propels the coat and the associated plasma membrane invagination, and covers the nascent endocytic vesicle after scission (Figure 1B and C). Lastly, vesicle scission was marked by the BAR-domain-containing Rvs167 proteins, with each species harboring one ortholog (Figure 1B and C) [34].

### Comparative analysis of endocytic protein assembly dynamics

To visualize and compare the endocytic assembly dynamics we tagged the proteins from the eight ortholog groups, with GFP at their C-terminus at the endogenous gene locus by homologous recombination. We imaged, by epifluorescence microscopy, live cells of each species that expressed these fusion proteins in identical conditions (EMM media at 24°C). All the proteins in the three species localized to small puncta at the plasma membrane consistent with a function in endocytosis (Figure 2A). The localizations were consistent with earlier reports for several *S. cerevisiae* and *S. pombe* proteins (*S.cerevisiae* proteins - Eps15, intersectin, Sla1, WASP, myosin Is, Rvs167 and fimbrin; and *S. pombe* proteins - intersectin, Sla1, WASP, myosin I, Rvs167 and fimbrin) [17,20,22,23]. Both *S. pombe* Eps15 paralogs were similarly localized, however, the Ede1 signal was brighter than the Ucp8 signal (Figure 2A).

**Figure 2:**
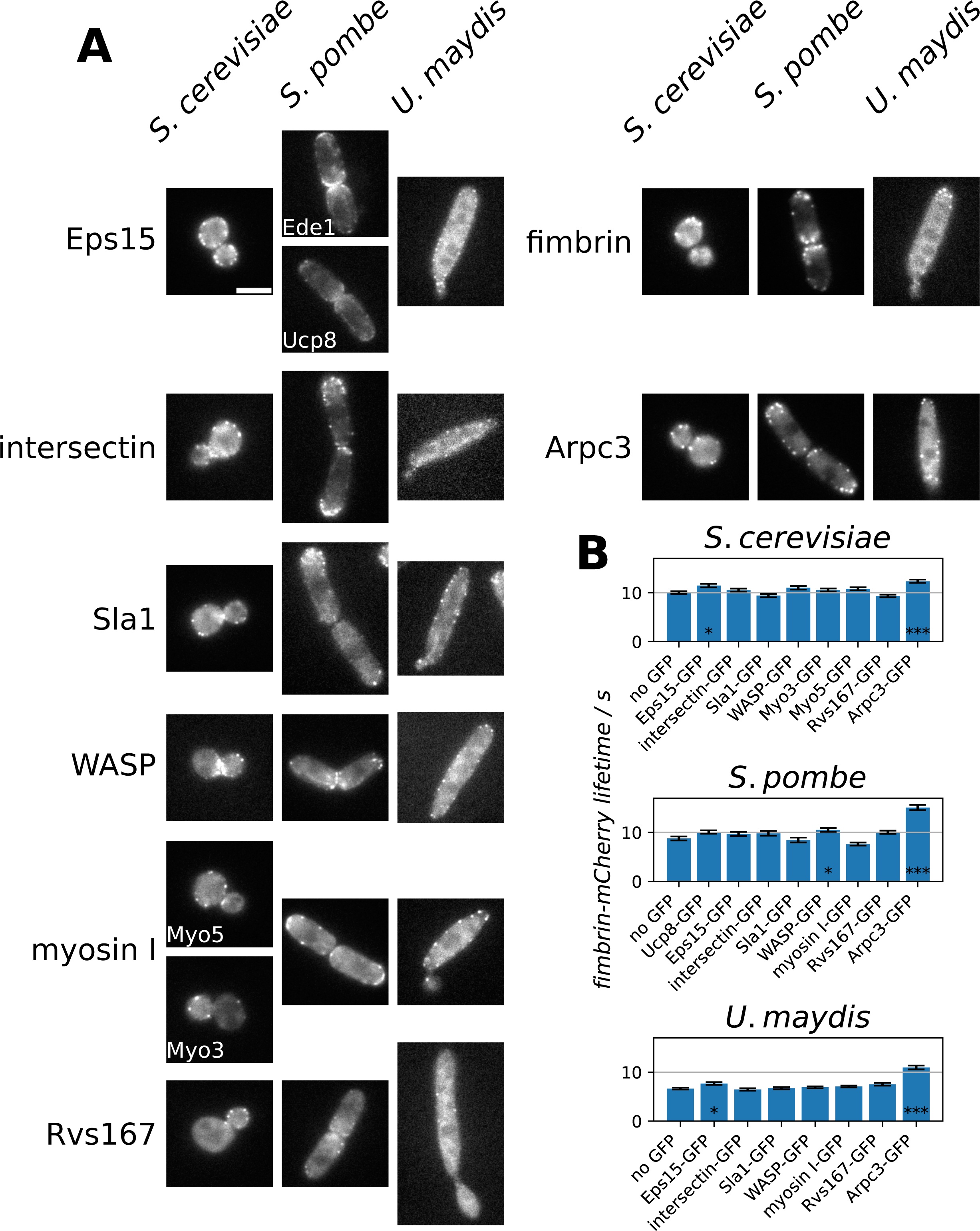
(A) Midplane epifluorescence images for different endocytic protein orthologs in each species. Scale = 5 μm. (B) A bar plot indicating fimbrin-mCherry lifetimes (mean ± S.E.) in cells when co-expressed with the indicated GFP-tagged orthologs.

We next compared the temporal organization of endocytic events. For each species, we generated strains that expressed each GFP-tagged endocytic protein in conjunction with the corresponding species’ fimbrin, endogenously tagged with mCherry. Fimbrin acted as a temporal marker for actin assembly. We also used the fimbrin patch lifetimes to ascertain functionality of the tagged endocytic proteins. We measured fimbrin-mCherry lifetimes when expressed either alone or together with each GFP-tagged ortholog. This is a sensitive approach to detect subtle perturbations in the endocytic protein functionality [21]. Most GFP-tagged orthologs minimally affected fimbrin lifetimes (Figure 2B). The tagged Arc3 orthologs, however, showed notably increased fimbrin lifetimes in all three species, which suggests that the GFP-tagging compromised Arc3 function (Figure 2B). We therefore excluded Arc3 from further analysis.

We then imaged the strains with live two-color total internal reflection fluorescence (TIRF) microscopy and we observed the appearance of each GFP-tagged protein in respect to the appearance of the actin marker, mCherry-tagged fimbrin (Figure 3). We used TIRF microscopy as it provides the best sensitivity for the often weak endocytic protein signals. Generally, the endocytic events progressed in a regular manner and terminated with actin assembly in all three species. However, lifetimes of the orthologs differed across species (Figure 3, Supplemental Figure S1).

**Figure 3:**
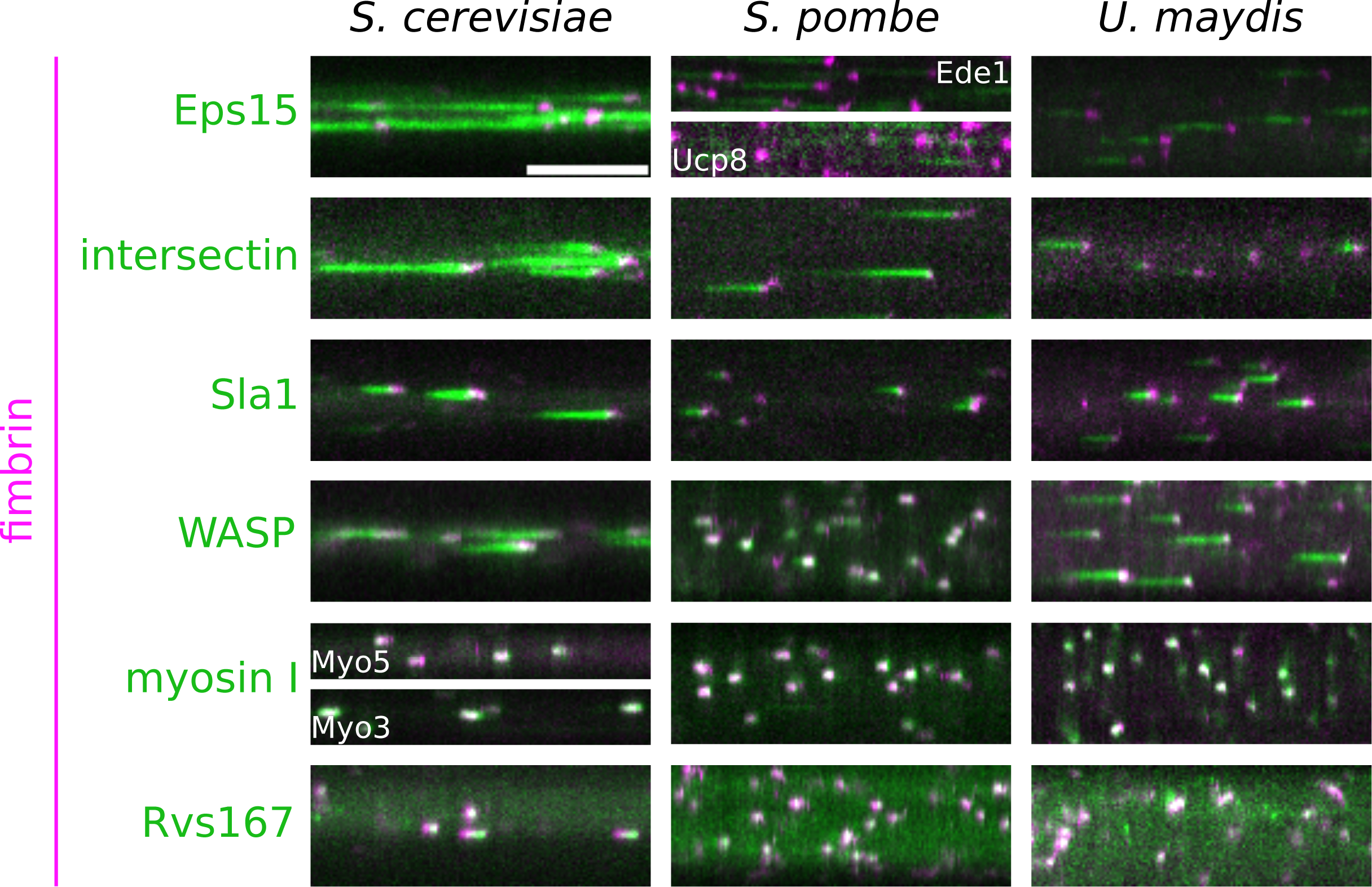
Kymograph representations of 2-color TIRF microscopy movies showing the indicated endocytosis orthologs, tagged with GFP (green), and the co-expressed fimbrin-mCherry (magenta), in each of the three species. Scale bar is 60 s.

To quantitatively compare the differences in the endocytic assembly dynamics in each species, we measured the lifetimes of each GFP-tagged ortholog, imaged together with the corresponding fimbrin-mCherry (Supplementary Table S1). We then registered these lifetimes in respect to the beginning of the invagination movement (Figure 4, see Materials and methods). These protein lifetime alignments represent the temporal choreography of the endocytic pathway of each species (Figure 4).

**Figure 4:**
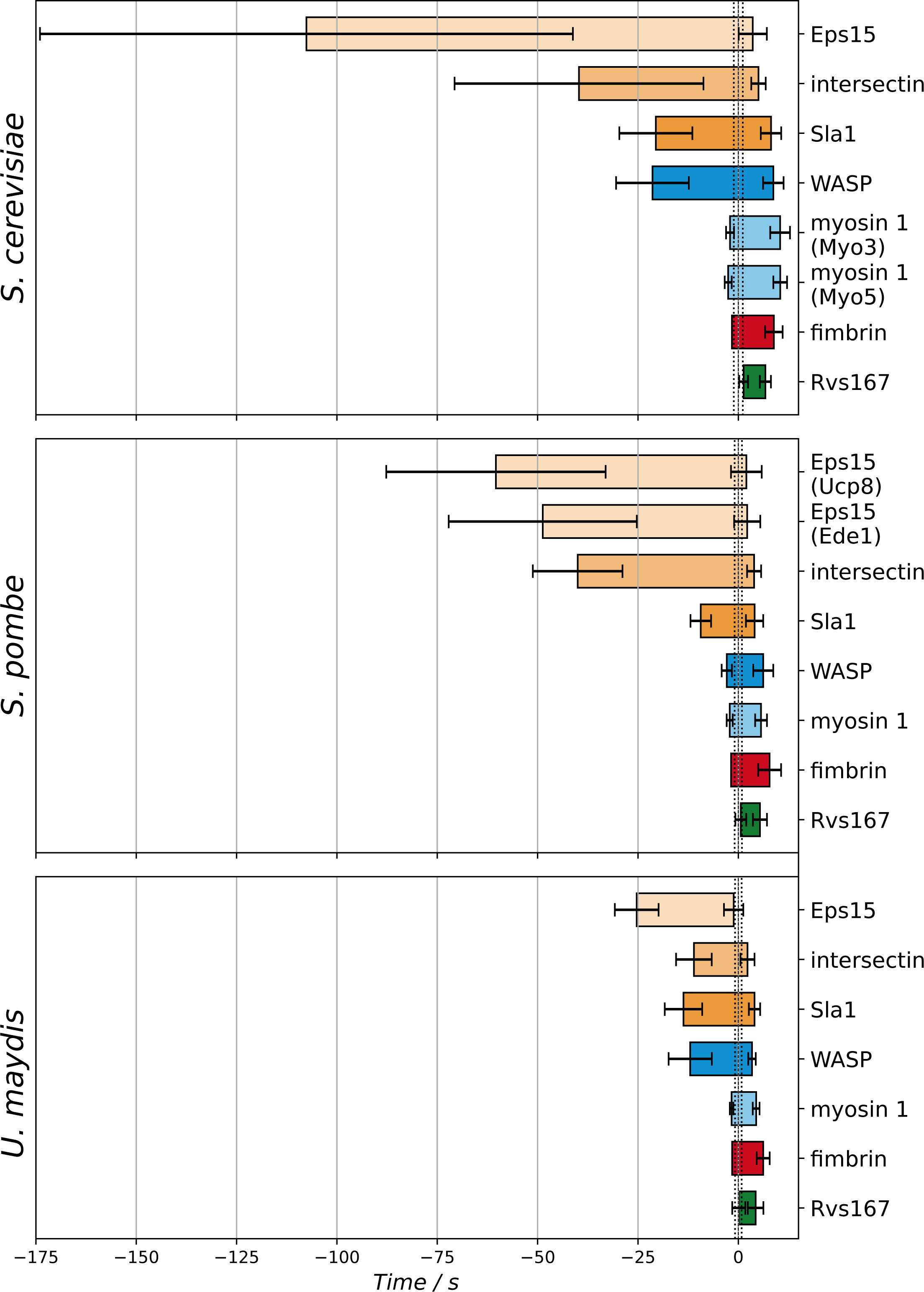
Bar graph showing the time of appearance and disappearance of each endocytic ortholog in respect to the beginning of the invagination movement, in each species. Error bars represent the SD for each measurement. Time 0 s marks the estimation of the beginning of the invagination movement. The dashed vertical lines represent its SD.

The entire endocytic process, from site initiation to disappearance of endocytic protein signal, was the longest in *S. cerevisiae*, spanning roughly two minutes (118 ± 66 s, mean ± SD). *U. maydis* endocytic process was the shortest, approximately half a minute long (32 ± 7 s, mean ± SD), while *S. pombe* endocytosis was intermediate, about one minute long (68 ± 27 s, mean ± SD). The Eps15 proteins arrived early in all species. *S. cerevisiae* Eps15 had the longest lifetime (111 ± 66 s, mean ± SD), followed by the two *S. pombe* orthologs (Ucp8 62 ± 28 s; Ede1 51 ± 24 s, mean ± SD). The lifetimes for *S. cerevisiae* Eps15 (coefficient of variation, CV: 0.59), and the two *S. pombe* Eps15s (Ucp8 CV: 0.45; Ede1 CV: 0.47) had a high variability (Figure 4 and Supplemental Figure S1). In contrast, *U. maydis* had a significantly shorter Eps15 lifetime (24 ± 6 s, mean ± SD) with significantly less variability compared to the two ascomycete species (CV: 0.25; Figure 4). The divergence in Eps15 dynamics suggests that the regulation of the endocytic early phase may have evolved differently in the ascomycetes (*S. cerevisiae* and *S. pombe)* and basidiomycetes (*U. maydis)*.

Intersectin, Sla1, and WASP function together in *S. cerevisiae* to regulate the localization and timing of actin assembly [35–37]. *S. cerevisiae* intersectin arrived slightly before Sla1 and WASP, consistent with previous observations [36]. Also in *S. pombe* intersectin preceded Sla1 and WASP, but both Sla1 and WASP had significantly shorter lifetimes than intersectin (Figure 4). The corresponding *U. maydis* proteins were shorter-lived than the orthologs in *S. cerevisiae*. In *U. maydis,* the assembly order was notably different: intersectin appeared slightly after Sla1 and WASP (Figure 4). Intersectin, Sla1 and WASP have critical roles in timing the actin assembly at endocytic sites [29,36,38]. Therefore these results reveal an unexpected variability in the temporal dynamics of critical endocytic regulators.

The late-arriving endocytosis proteins myosin I, Rvs167, and fimbrin had short lifetimes and appeared similar across species (Figure 4). The myosin I orthologs arrived when actin assembly began in all three species. However, the two myosin I orthologs in *S. cerevisiae* were slightly longer-lived compared to the other two yeasts, which indicates different myosin disassembly dynamics after vesicle budding.

These results show that the three fungal species use a sequential assembly from early to late endocytic proteins. However, the overall duration of the endocytic process differs between species. Particularly the early phase of endocytosis is different in duration. Furthermore, the assembly timing and even the assembly order of endocytic proteins is species specific. Thus, the lifetimes of individual protein orthologs can evolve in a mosaic manner, independent of the total duration of endocytosis.

### Comparative analysis of endocytic protein motility

In addition to having a precise sequential assembly, the endocytic proteins exhibit different motile behaviors that are indicative of where they are located at the endocytic site during vesicle budding [20]. The movements of endocytic proteins can be measured by centroid tracking of endocytic patches imaged at the midplane of living cells [21,23,39,40]. We used the centroid trajectories of coat and scission proteins to estimate the length and growth rate of the endocytic membrane invagination, and the time of vesicle scission [21]. We used this approach to investigate how endocytic events differ in their spatial dynamics in the different fungi. First, we imaged Sla1-GFP orthologs, which show the movement of the endocytic coat and serve as a proxy for the growth of the endocytic membrane invagination. Sla1 orthologs showed qualitatively similar behavior in each species by first assembling into a stationary patch, which later moved into the cell and then rapidly disassembled (Figure 5A). We then performed centroid tracking analysis on Sla1-GFP in each species and aligned the average centroid tracks in time to the moment when the Sla1 movement starts (Figure 5B, upper panel). In each species, Sla1 centroids moved a different distance into the cell. In *S. pombe*, Sla1 internalized the furthest (∼300 nm), followed by *S. cerevisiae* (∼150 nm) and lastly *U. maydis*, which only internalized ∼100 nm. Surprisingly, during the first 100 nm of movement all the Sla1 orthologs internalized at nearly identical rates (Figure 5B). The assembly timing of Sla1, indicated by the fluorescent intensity profiles, varied across the species. However, the intensities all peaked near the beginning of the coat movement, which likely reflects the critical actin regulatory role of Sla1 (Figure 5B, lower panel).

**Figure 5:**
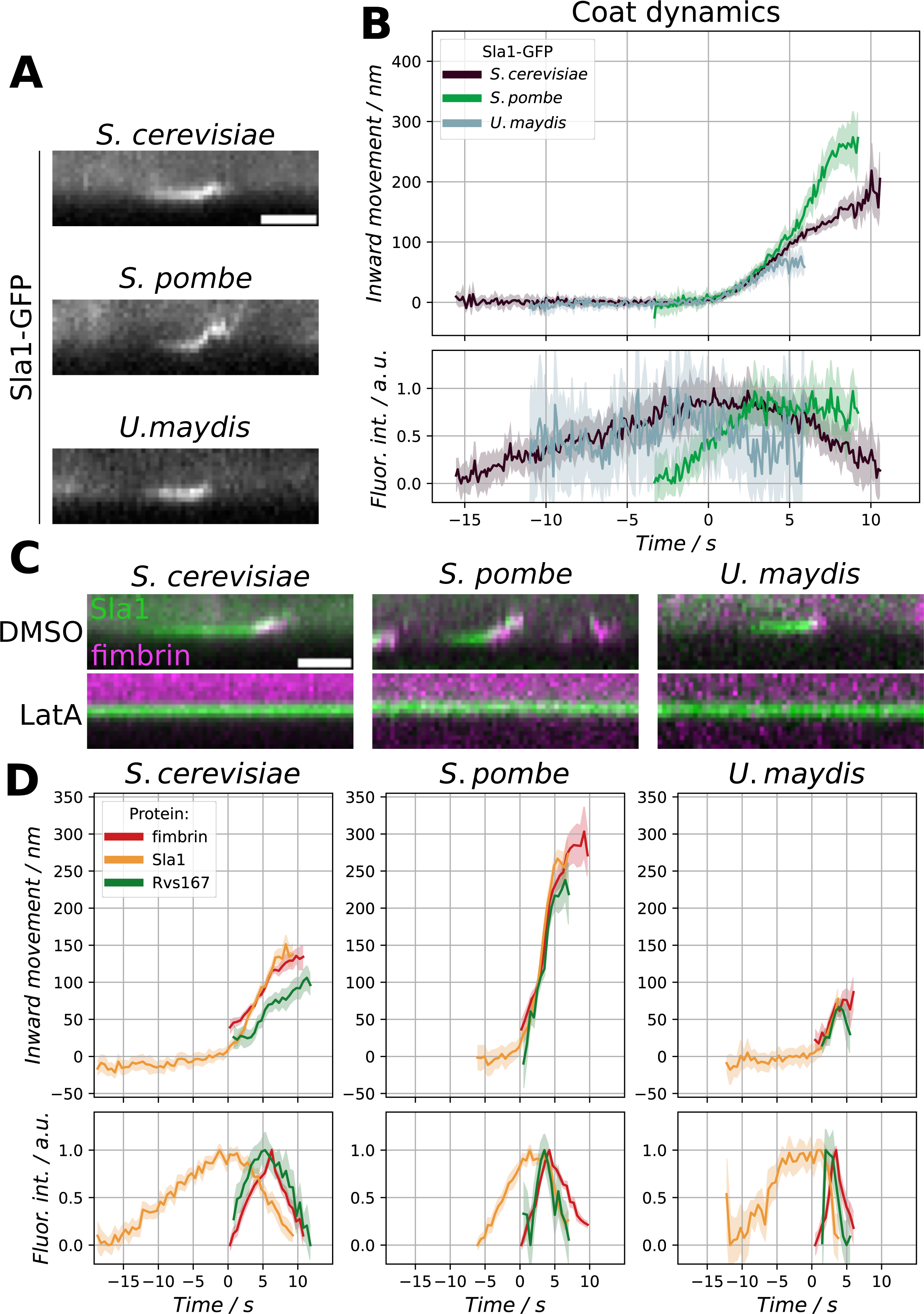
(A) Kymograph representation of Sla1-GFP ortholog dynamics imaged with mid-plane epifluorescence. The kymograph is oriented so that the cell outside is at its base while the cell cytoplasm at the top. Scale bar is 30 s. (B) Average trajectories of Sla1-GFP dynamics depict the coat inward movement during endocytosis and the fluorescence intensity changes in the endocytic patches due to Sla1-GFP accumulation and disassembly. The shading represents the 95% CI (C) Kymograph representation of Sla1-GFP and Fim1-mCherry ortholog dynamics imaged with mid-plane epifluorescence, in cells treated with DMSO or latA. Scale bar is 30 s. (D) Average internalization dynamics of Sla1, Rvs167, and fimbrin during the endocytosis, and their average change in fluorescence intensity. The shading represents the 95% CI

The internalization movement and coat disassembly is known to be strictly actin-dependent in *S. cerevisiae* and *S. pombe* [20,41]. We treated cells that expressed Sla1-GFP and fimbrin-mCherry orthologues with latrunculin A, which inhibits actin polymerization. This treatment prevented the formation of fimbrin-mCherry patches in each species, and showed that Sla1 internalization and disassembly were actin-dependent in *U. maydis*, as in *S. cerevisiae* and *S. pombe* (Figure 5C).

The time of vesicle scission in *S. cerevisiae* is marked by the peak of fluorescence intensity and rapid centroid movement of Rvs167-GFP [21,28]. We tracked Rvs167-GFP and fimbrin-mCherry dynamics to time the scission event in respect to fimbrin. We then tracked Sla1-GFP and fimbrin-mCherry to register the scission time in respect to the coat internalization movement (Figure 5D, see Materials and methods). The distance that Sla1 centroids move until the scission time gives an approximation of the invagination length [21]. These data suggest that the membrane invagination length at the time of scission is different in each species: ∼100 nm in *S. cerevisiae*, ∼150 nm in *S. pombe*, and ∼50 nm in *U. maydis.* Furthermore, we can conclude that relative to the start of invagination the scission events are similarly timed in the ascomycete species, while in *U. maydis* scission occurs earlier.

In summary, our findings show that endocytic membrane invagination in each species is strictly actin-dependent and happens with a similar initial rate. However, the progression of membrane invagination and the timing of vesicle scission have changed during evolution.

### Robustness of endocytic dynamics against environmental variation

Our results suggest an evolutionary divergence in the dynamics of endocytic protein assembly and vesicle budding between the three species. However, our results were obtained under a single set of environmental conditions, and it is possible that the endocytic process has been optimized to different environmental conditions in these species. Therefore, the observed differences could simply be a function of the environmental conditions used in the experiments. To test this idea, we chose temperature as an environmental variable, and performed the Sla1-GFP centroid tracking analysis at five different temperatures: 18, 21, 24, 27 and 30°C, which all support growth of each species. The Sla1-GFP patches were assembled and internalized at all tested temperatures, which indicates that endocytosis is functional (Figure 6A and Supplemental Figure S2). However, the Sla1 behavior in the three species was differently impacted by temperature (Figure 6A and Supplemental Figure S2). We quantified three parameters from the Sla1 tracking data; (1) the average lifetime of Sla1 patches, as a measure of the duration of the late phase of endocytosis, (2) the velocity of the centroid movement during the first 50 nm to estimate the growth speed of the early invaginations, and (3) the distance that the centroid is internalized, as a proxy for the maximal length of the membrane invaginations (Figure 6B, C and D). Sla1 lifetimes for each species shortened with temperature increase, which suggests that the overall progression of endocytosis is sped up (Figure 6B). The internalization velocity changed at different temperatures for each species (Figure 6C). The velocities of *S. cerevisiae* and *S. pombe* Sla1 patches increased with increase in temperature while *U. maydis* showed almost no change over the temperature range. Unlike the other two parameters, the Sla1 centroid displacements were strikingly similar over the temperature range suggesting that the endocytic invagination length is robustly buffered against temperature variations (Figure 6D).

**Figure 6:**
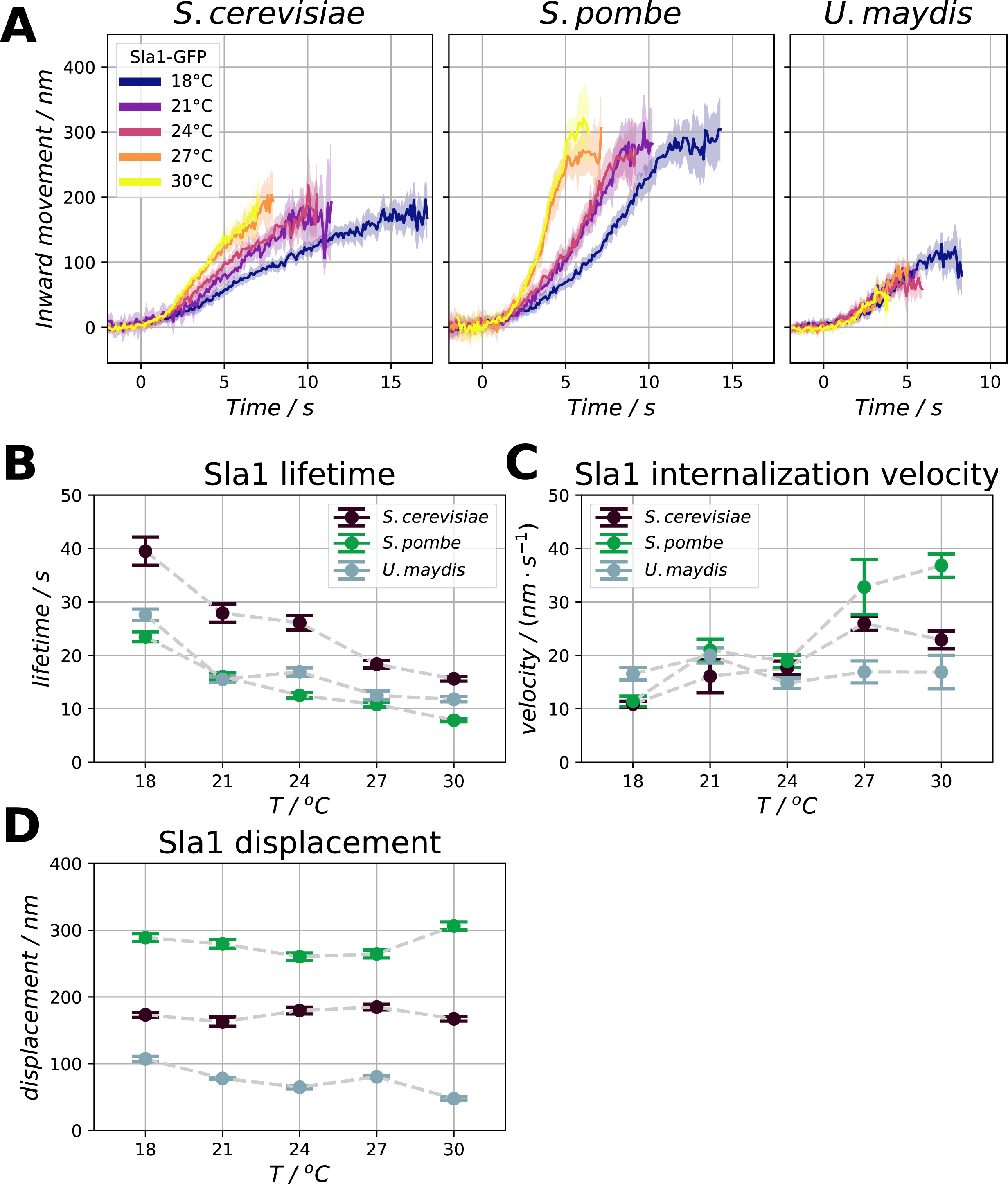
(A) Inward movement of a coat marker, Sla1-GFP, for the three species at 18°C, 21°C, 24°C, 27°C, and 30°C. The shading represents the 95% CI (B, C, and D) Reaction norm curves of the three species for the coat marker Sla1-GFP lifetime (B), the magnitude of its inward movement (C), and its speed of internalization (D). Error bars represent the standard error.

Taken together these results show that endocytosis exhibits a complex response to temperature, with some parameters being phenotypically plastic, while others, such as the invagination length, being robustly buffered. The invagination length thus appears to be a species-specific, evolved phenotypic trait and not simply a differential response to the environment.

### Evolution of WASP assembly

The function of WASP has been studied extensively in both *S. cerevisiae* and *S. pombe* [29,38]. At the endocytic site WASP binds to and activates the Arp2/3 complex to nucleate new actin filaments [29]. In *S. cerevisiae*, WASP molecules are assembled at the endocytic site before the actin assembly starts, while in *S. pombe* assembly of WASP and actin happens almost simultaneously [23,39]. In *S. cerevisiae* the early assembly of a WASP ring around the endocytic coat is thought to ensure even distribution of actin-generated forces during coat internalization [42]. Thus this interspecies variation in the assembly of WASP (Figure 4) provided an attractive opportunity to investigate the molecular mechanism behind an evolutionary change. We measured the assembly dynamics of WASP-GFP orthologs co-expressed with the actin marker, fimbrin-mCherry. This analysis showed that in *U. maydis* and *S. cerevisiae* most of the WASP molecules were assembled at the endocytic site before the actin assembly started (Figure 7A). We refer to this assembly phenotype as WASP pre-assembly. In *S. pombe,* WASP and actin assembly occurred nearly simultaneously (Figure 7A), which we defined as a co-assembly phenotype.

**Figure 7:**
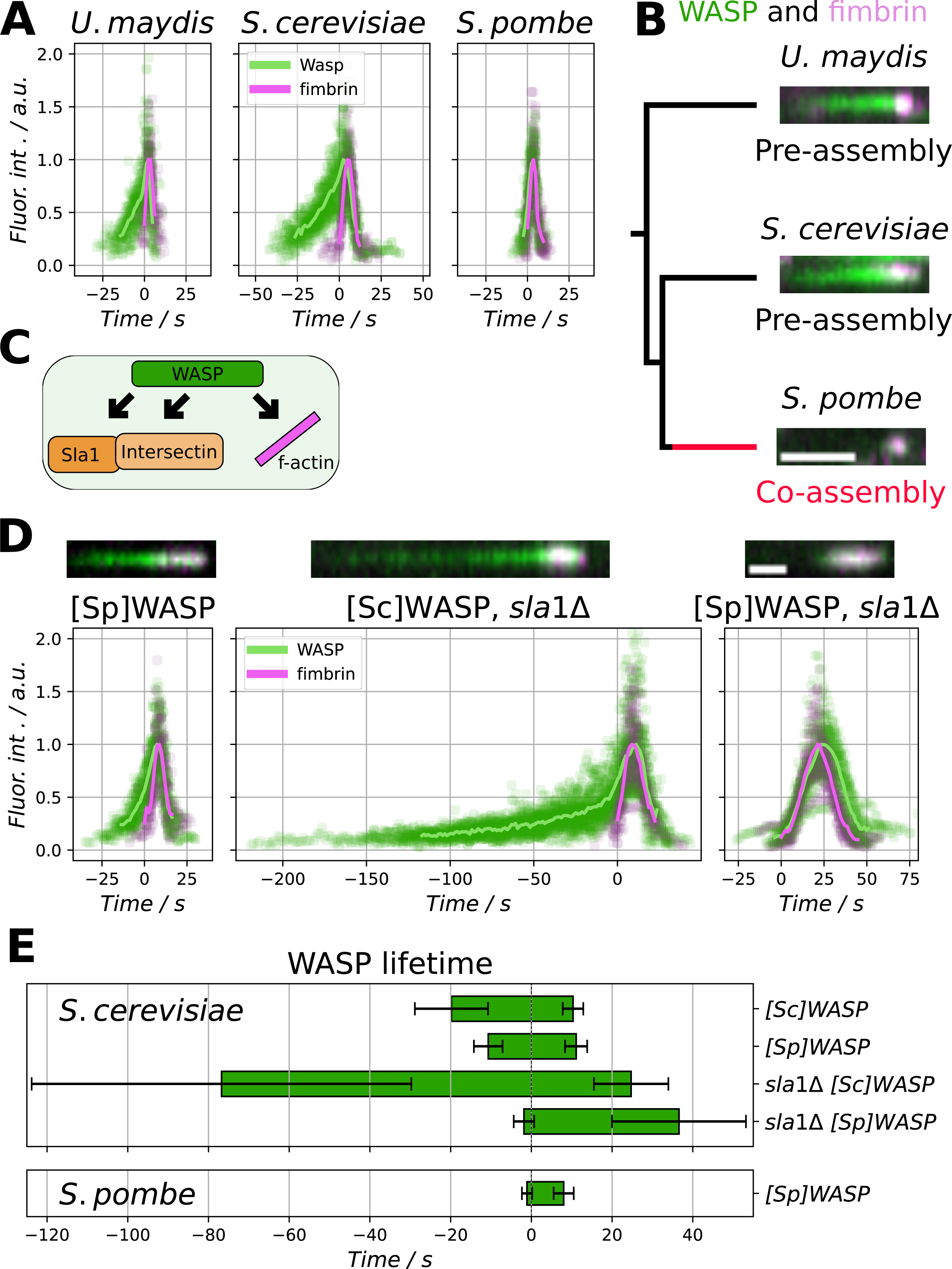
(A) Individual data points of fluorescence intensity curves and their average for WASP and fimbrin in *S. cerevisiae, S. pombe*, and *U. maydis.* The time point 0 marks the appearance of fimbrin. (B) Kymograph representation of WASP and fimbrin in the three species. The kymographs are taken from TIRF movies. The kymographs are related by an evolutionary tree. In *S. cerevisiae*, and *U. maydis* WASP pre-assembles before fimbrin, while in *S. pombe* WASP and fimbrin co-assemble. Scale bar marks 20 s. (C) Sla1 and intersectin regulate WASP activity; WASP promotes actin filament nucleation, and the actin network promotes the recruitment of WASP. (D) Left panel: kymograph representations of [Sp]WASP and fimbrin appearance in *S. cerevisiae*, and their fluorescence intensity curves, shown as individual data points of fluorescence intensity curves and their average. Middle panel: kymograph representations of [Sc]WASP and fimbrin appearance in *S. cerevisiae* strains missing Sla1, and their fluorescence intensity curves. Right panel: kymograph representations of [Sp]WASP and fimbrin appearance in *S. cerevisiae* strains missing Sla1, and their fluorescence intensity curves. The kymographs are taken from TIRF movies. Scale bar marks 10 s. (E) [Sc]WASP and [Sp]WASP lifetimes in *S. cerevisiae* cells with and without Sla1, and [Sp]WASP lifetime in *S. pombe* cells shown for comparison. Error bars mark the SD.

We used the phylogenetic relationships of the species to estimate the likely direction of evolutionary change of the pre-assembly and co-assembly phenotypes (Figure 7B). The WASP assembly differs between *S. cerevisiae* and *S. pombe*, which are phylogenetically closer than *U. maydis*, but this species shares a WASP pre-assembly with *S. cerevisiae*. Therefore, pre-assembly is likely the ancestral state and the co-assembly phenotype of *S. pombe* an evolutionary novelty (Figure 7B).

In *S. cerevisiae* intersectin and Sla1 recruit WASP to the endocytic site [36]. In addition, *S. cerevisiae* WASP binds to actin filaments *in vitro*, which may also facilitate its recruitment [43](Figure 7C). In *S. pombe,* the direct binding of WASP to actin filaments is proposed to contribute to WASP recruitment [44,45], while an intersectin and Sla1 role is not established (Figure 7C). Changes in these recruitment mechanisms may influence pre-assembly to co-assembly phenotypes.

We hypothesized that the WASP sequences themselves could contain the information about the assembly mode. We tested this idea by replacing the *S. cerevisiae* WASP ([Sc]WASP) coding region with the homologous *S. pombe* WASP ([Sp]WASP). We expressed [Sp]WASP as a GFP fusion in *S. cerevisiae* with fimbrin-mCherry and measured the assembly dynamics (Figure 7D). In *S. cerevisiae,* [Sp]WASP exhibited a pre-assembly behavior, although the preassembly period was slightly shorter than for the endogenous [Sc]WASP (Figure 7E). This result suggests that key determinants of the WASP assembly timing are present in other proteins of the endocytic machinery.

Next, we aimed to promote WASP co-assembly phenotype in *S. cerevisiae* by weakening the potential pre-assembly pathway by deletion of Sla1 [36]. In the absence of Sla1, [Sc]WASP showed a biphasic assembly pattern. First, WASP was pre-recruited at a very slow rate. Then, when actin started assembling, WASP was recruited at a more rapid rate that resembled the dynamics of WASP co-assembly (Figure 7D, 7E; [35]). We hypothesize that these two WASP assembly phases may represent two mechanisms: a slow, defective pre-recruitment followed by a fast co-recruitment driven by binding to actin filaments.

We then combined the deletion of Sla1 with [Sp]WASP. This strain exhibited a complete loss of WASP pre-assembly and retained only a striking co-assembly of WASP with actin (Fig 7D). These results reveal that the WASP assembly timing is encoded both in WASP itself and in the proteins recruiting it to the endocytic site. The lifetime of fimbrin patches was increased in the *S. cerevisiae* strain expressing [Sp]WASP and harboring Sla1 deletion (Fig 7D), indicating that the genetic manipulation is not entirely neutral for the normal progression of endocytosis.

Taken together these results indicate that relatively few genetic changes are needed to shift the WASP assembly phenotype. Furthermore, they allow us to suggest a possible evolutionary pathway for the change in WASP assembly. The likely ancestral state was WASP pre-assembly mediated by Sla1 and Pan1. The preassembly ancestral state was maintained in *S. cerevisiae* and *U. maydis*. The co-assembly likely evolved in the *S. pombe* lineage by weakening of intersectin and Sla1-mediated WASP recruitment leaving the actin-mediated co-assembly as the dominant mechanism.

## DISCUSSION

In this study we compared quantitatively the dynamic phenotypic traits of clathrin-mediated endocytosis in three species within the dikarya clade of fungi. These three fungal species are separated by more than 500 million years of evolution, but share a similar overall cellular architecture and lifestyle. We found that the overall modular and sequential progression of endocytosis was conserved across the three species, but there was a mosaic of differences in the assembly sequence and in vesicle formation, which suggested that the endocytic mechanisms have evolved.

### Evolution of endocytic coat assembly

The average total duration of an endocytic event varied fourfold between the species (Figure 8A). This difference was largely accounted for by the endocytic early phase, which varied in duration and regularity between species. Studies in *S. cerevisiae,* have suggested that endocytic cargo sorting predominantly occurs during the early phase [46]. The early phase starts by the assembly of Eps15, clathrin, and several adaptor proteins, and ends with the arrival of the late coat proteins intersectin, Sla1, and End3 [47]. The duration of the early phase has been proposed to depend on the cargo recruitment rate, so that there would be a “cargo checkpoint” that controls the transition from the early to late phase [47,48]. The molecular mechanism of the cargo checkpoint remains unknown, but there is evidence from *S. cerevisiae* and mammalian cells to support the idea [49–51]. The early phase of *U. maydis* is unexpectedly short and regular relative to the ascomycetes. This may indicate that either the rate at which the endocytic sites are filled with cargo is faster in *U. maydis*, or that the transition from the early to late phase is not controlled by a cargo checkpoint in this species. The rate at which the full complement of cargo is recruited could be influenced by the concentration of available cargo on the plasma membrane and the cargo capacity of the subsequent vesicle. The early phase variability could reflect different priorities for endocytic uptake such as specific cargo transport versus recycling of the plasma membrane lipids.

**Figure 8:**
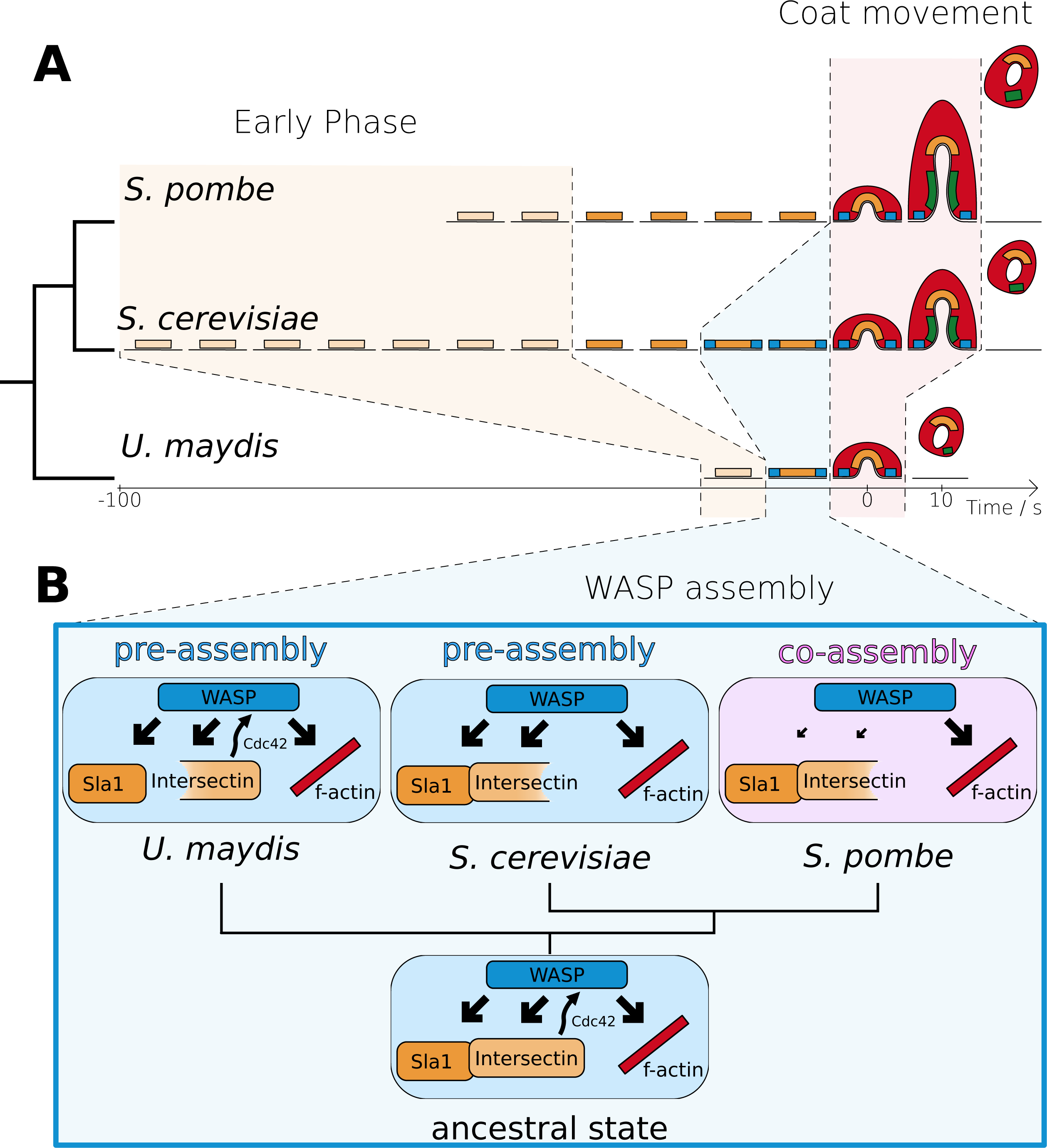
(A) Representation of the different module dynamics in *S.cerevisiae*, *S. pombe*, and *U. maydis.* The coloured areas highlight the “Early phase”, the “WASP assembly phase”, and the “Coat movement”, which all show distinct dynamics in the three species. (B) Some of the molecular interactions that can be responsible for the differences in the pre-assembly and co-assembly mechanisms, and their possible ancestral state.

The assembly of the late coat proteins also showed unexpected variation. In *S. cerevisiae,* intersectin interacts directly with Sla1 and recruits it to the endocytic sites [36,52]. Consistently, in *S. pombe*, the arrival of intersectin precedes Sla1. However, the assembly order is reversed in *U. maydis*. This difference suggests that the assembly mechanism for the late coat proteins has diverged between the two fungal lineages of ascomycota and basidiomycota. Interestingly, in animals, intersectin is among the first proteins to arrive at the endocytic site [53], which suggests that animal cells lack distinct early and late phases as defined for fungi. Therefore, the separation of early and late phases may be an evolutionary innovation in fungi.

### Evolution of WASP assembly and regulation

The key regulator of actin patch assembly, WASP, showed surprising variability in its assembly dynamics that offered an opportunity to investigate the mechanistic basis of phenotypic divergence. We distinguished two diverged WASP assembly phenotypes: an ancestral pre-assembly of WASP before actin, in *S. cerevisiae* and *U. maydis*, and an evolutionarily novel co-assembly of WASP together with actin, in *S. pombe*. The pre-assembly of WASP at the rim of the endocytic coat was proposed to be important for isotropic nucleation of actin filaments at the endocytic site and thereby for even distribution of actin-derived forces to efficiently shape the membrane invagination [42]. The co-assembly phenotype in *S. pombe* indicates that either WASP pre-assembly is not functionally relevant as proposed or that *S. pombe* has evolved another mechanism to guarantee isotropic actin nucleation. The clathrin-mediated endocytosis of mammalian cells and *Dictyostelium discoideum* exhibit a WASP co-assembly phenotype [30,54]. This implies that co-assembly may be the ancestral state in the supergroup amorphea, which contains fungi, animals and amoebozoa. WASP pre-assembly may be an evolutionary novelty in fungi, and possibly linked to increased force requirements in membrane shaping in fungi compared to animals [7]. Intriguingly, *S. pombe* has evolved a WASP co-assembly that resembles animal and amoeba co-assembly states.

In *S.cerevisiae* intersectin and Sla1 are the key WASP recruiters for pre-assembly [36]. Conversely, the co-assembly in *S. pombe* may largely depend on positive feedback based on the ability of WASP to be recruited by binding directly to actin filaments [44,45]. The actin filament-binding activity of WASP is conserved in *S. cerevisiae* [43], which suggests that the co-assembly capacity was already encoded in the ancestral endocytic machinery. The co-assembly phenotype, however, may not be expressed as long as WASP is efficiently pre-recruited before actin filament assembly. Weakening of the pre-assembly mechanism by a deletion of *SLA1* caused *S. cerevisiae’s* endogenous WASP to exhibit a partial co-assembly phenotype: WASP first pre-assembled at a slow rate and then rapidly peaked together with actin patch assembly. We were able to completely change WASP pre-assembly into a co-assembly phenotype in *S. cerevisiae* by replacing the endogenous *S. cerevisiae* WASP with the *S. pombe* WASP in *-./”0* cells. The idea that the co-assembly phenotype evolved based on pre-existing biochemical activities may explain how we were able to change the WASP assembly phenotype by simple genetic modifications of two genes. This result also shows that gene swapping experiments can provide a way to uncover the molecular changes that have led to cellular phenotype evolution.

### Evolution of vesicle budding

In all three species the membrane invagination is actin-dependent and takes place within the final 10s of the endocytic process. This is drastically different from animals where membrane invagination starts early during endocytosis and is only partially actin-dependent [55,56]. The coat internalization rate at 24°C was virtually identical in the three species for the first ∼100 nm (Figure 5). However, after 100 nm the species differed significantly. *S. cerevisiae* coat moves up to ∼200 nm and *S. pombe* to almost 300 nm. Despite these length differences, in both *S.pombe* and *S. cerevisiae* Rvs167 scission occurs about 8 s after invagination initiation. *U. maydis* stops its coat movement at ∼100 nm and the scission takes place earlier, only about 4 s after invagination starts (Figure 5). These observations suggest that the endocytic invaginations are of different maximum length in different species and may yield vesicles of different sizes. Mutant studies in *S. cerevisiae* have suggested that invagination length correlates with the vesicles size [28]. Similar initial invagination rates suggest that the actin and myosin based molecular mechanisms driving invagination growth are conserved. The maximal length of the invagination may be modulated by evolutionary changes in the mechanisms that determine when actin nucleation is turned off and when the vesicle scission happens.

### Molecular clues to phenotypic variation

Sequence level divergences in endocytic proteins provide some clues about the molecular mechanisms that underlie the phenotypic evolution. *U. maydis* intersectin and WASP contain RhoGEF and CRIB domains, respectively, similar to the mammalian orthologs [26,57] (Figure 1C and 8B). In mammalian cells and in basidiomycetes, WASP is regulated by conformational changes induced by the CRIB domain interaction with the GTPase Cdc42 and intersectin has reported GEF activity on Cdc42 [58–61]. Loss of this additional WASP regulation may have opened new evolutionary paths for WASP assembly within the ascomycetes. In addition, *U. maydis* intersectin lacks the first EH domain (Figure 1 and 8B), which binds to and recruits Sla1 in *S.cerevisiae* [36]. *U. maydis* also lacks a detectable End3 ortholog, which participates in Sla1 recruitment in *S.cerevisiae* [36]. The loss of these Sla1 interactions may contribute to the assembly order differences of intersectin and Sla1 in *U. maydis* (Figure 4). Furthermore, the numbers of endocytic protein molecules at the endocytic site differ between species. *S. pombe* has more actin at endocytic sites than *S. cerevisiae* [17,23]. Actin patch size and different endocytic protein stoichiometries may affect the endocytic phenotypes, such as for the invagination length and WASP patterning for isotropic actin nucleation.

### Effect of the environment

Phenotypes are expressions of both the genotype and the environment. Evolutionary adaptation to a specific environment, such as a certain temperature range, may lead to reduced performance under different environmental conditions [62]. This is an important consideration when comparing phenotypic traits in different species under standardized conditions. To address environmental contributions we analyzed the effect of temperature. In physiological temperature ranges, many cell biological processes, including protein binding and membrane fluidity, follow Arrhenius’ law, which describes how reaction rates depend on temperature [63].

We used a temperature range which supports growth of all three species, and measured three phenotypic parameters: lifetime of the late coat, invagination growth rate and coat displacement. The Sla1 lifetimes steadily decreased as temperature increased, which indicates a faster endocytic process. This result is consistent with Arrhenius’ law and earlier observations that mammalian clathrin-mediated endocytic uptake correlates with temperature, which is likely due to changes in membrane fluidity and protein binding rates [64,65]. However, we found that invagination rates change only at the upper range of the tested temperatures, and coat displacements are invariant in this temperature range. This suggests that cells actively maintain an optimum over a range of temperatures or that these traits are based on temperature independent parameters, such as endocytic protein amounts or structural features of the endocytic machinery. The molecular mechanisms of endocytosis that respond to environmental parameters, and the evolutionary pathways that regulate these responses are a little studied area and fertile ground for future studies.

### Future directions

Our results demonstrate that comparative cell biology provides new mechanistic insights into evolutionary change of clathrin-mediated endocytosis, a core cellular pathway. This approach requires quantitative data from several species, which are relevant to the evolutionary question asked. Obtaining such data from multiple species remains a big challenge. However, recent advancements in genome sequencing, genetic manipulation techniques, protein structure determination and prediction, and emergence of new model organisms are removing these technical barriers. Merging evolutionary and cell biology will be critical to uncover the evolutionary origins of cellular processes, link the molecular evolution to evolution of cellular phenotypes, and clarify the role of adaptation versus random genetic drift that drives cell level evolution [66].

## MATERIALS AND METHODS

### Strains and media

Strains were generated in the following backgrounds MKY100 (this group), YSM1371 (Sophie Martin lab) and UM521 (Fungal genetics stock center) for *S. cerevisiae*, *S. pombe* and *U. maydis*, respectively. *S. cerevisiae* were grown in rich medium (YPD) or synthetic medium (SD) and supplemented with appropriate amino acids or antibiotics. *S. pombe* were grown in rich medium (YES) or synthetic medium (EMM) and supplemented with appropriate antibiotics. *U. maydis* were grown in rich medium (YEPS-light) or synthetic medium (EMM) and supplemented with appropriate antibiotics. All strains were cultured at 24°C unless otherwise specified. All media formulations and antibiotics are listed in Supplemental Table S2.

C-terminal eGFP (*S. cerevisiae* and *U. maydis*), superfold GFP (*S. pombe*) or mCherry tags were integrated chromosomally to generate C-terminal fusions of each protein. Gene construction and transformation protocols were performed as previously described for each species [67–69]. All tagging constructs were verified by PCR and sequencing. Original plasmids used and sources are listed in Supplemental Table S3. All strains used in this study are listed in Supplemental Table S4.

### Microscopy setup

All data were acquired on an IX81 and IX83 Olympus wide-field epifluorescence microscopes. The two microscopes were equipped as follows:IX81 Olympus microscope: a 488 nm and a 561 nm laser, an OBS-U-M2TIR 488/561 (Semrock, Rochester, NY) dichroic mirror, a CherryTemp system to control the temperature, an optional DualView (Optical Insights, LLC, Tucson, AZ) beam splitter, and a Hamamatsu ImagEM camera (C9100-13). The microscope was controlled with Micromanager (v2.0.1). The CherryTemp system was controlled with the CherryTemp software v1.0.1.45.

IX83 Olympus microscope: a 488 nm and a 561 nm laser, an Ilas2 system for TIRF, a TRF89902-OL3 (C174893) quad-filter cube, a Hamamatsu ImagEMx2 camera (C9100-23B), and an autofocus. The microscope was controlled with VisiView (v4.4.0.11, Visitron Systems GmbH).

### Phylogenic analysis

Orthologies were mined with PSI-BLAST and blastp at https://www.ncbi.nlm.nih.gov/. Reciprocal BLAST searches were done to confirm orthology.

### Sample preparation for imaging

First, cells were grown overnight at the imaging temperature (18°C, 21°C, 24°C, 27°C, or 30°C). Sp and Um were grown in EMM; Sc were grown in SC-Trp. Then, all cells were resuspended in EMM at the imaging temperature to the early logarithmic phase (OD_600: 0.4-0.5). When not stated, the imaging temperature was set at 24°C. To standardize imaging conditions, the three yeast species carrying fluorescent markers for the same orthologs were mixed together in the same sample. Cells were adhered on 25mm round coverslips (Menzel-Gläser, #1) by coating the coverslip with Concanavalin IV (ConA, 4mg/ml, Sigma-Aldrich C2010-100MG) by incubating the coverslip with ConA for 20 min, washing the excess of ConA, and then incubating the cells for 20 min. A last round of washing was necessary to remove cells that adhered poorly.

### Imaging

To compute protein lifetimes in Fig. 4 and 7 and to picture all the kymographs in Fig. 3 and 7, cells were imaged in TIRF mode on the IX83 Olympus microscope. Exposure times were set to 500 ms for the GFP and 200 ms for the mCherry. The EM-CCD gain was always at 255. Laser intensities were adjusted to 1-2% for the 488 nm laser and 15-30% for the 561 nm laser.

The beginning of the invagination movement was resolved by alternating epifluorescence and TIRF illumination. Laser intensities were set to 2% for the 488 nm laser and 25% for the 561 nm laser, and the exposure time was 200 ms for both channels. The beginning of the invagination movement was determined from the ratio of the two images (see Image and data analysis).

All other images and the images to track endocytic dynamics were acquired with the IX81 Olympus wide-field epifluorescence microscope. Data for Fig. 5A-B, 6 and Sup. Fig. 2 were acquired by imaging only the GFP channel, with an exposure time set to 100 ms, 32% 488 nm laser power, and EMCCD gain set to 255. We used a small ROI of 192 x 192 pixels to minimize the lag between subsequent frame acquisitions. The images for the data in Fig.5C-D were acquired simultaneously on the GFP and mCherry channels with the DualView beam splitter. The exposure time was 250ms, with 25% of 488 nm laser power and 63% of 561 nm. EMCCD gain was set to 255. The cells of the different species weren’t mixed together only to simultaneously acquire the GFP and mCherry channels. We used a small ROI of 416 x 128 pixels to minimize the lag between subsequent frame acquisitions. The ROI was large enough to encompass the beam splitter projection of the two channels on the sensor. The two channels were cropped from the ROI and registered together based on the two-color acquisition of Tetraspec beads (Picco and Kaksonen, 2017).

### Image repository

The images acquired and used in this work are published in the Zenodo repository:

10.5281/zenodo.10879761 (https://zenodo.org/records/10879761)

### Image and data analysis

All images were background subtracted and corrected for photobleaching in Fiji/Image (Picco and Kaksonen, 2017 Methods in Cell Biology).

Lifetimes were computed manually by observing the appearance and disappearance of GFP and RFP spots in the time series. To align lifetimes within species, the appearance of fimbrin-RFP served as a fiducial to align all the different protein lifetimes to the appearance of the actin network. To align lifetimes across species, the beginning of the invagination movement was estimated from the ratio between subsequent epifluorescent and TIRF acquisitions of an internalizing coat marker (Sla1-GFP). It was timed with respect to the appearance of fimbrin-mCherry, which served as a marker to register the beginning of the invagination movement with respect to all the other protein lifetimes.

Trajectories in Figure 5D were treated in pairs: Sla1-GFP and fimbrin-mCherry or Rvs167-GFP and fimbrin-mCherry. The pairs were generated by tracking the green and red patches in each endocytic event imaged simultaneously with the DualView beam splitter. Each pair was aligned to an average trajectory for fimbrin-GFP, which we generated *a priori*, imaging fimbrin-GFP alone (as for Sla1-GFP alone). The alignment was done by aligning in time each fimbrin-mCherry to the average trajectory for fimbrin-GFP (‘align_raw’ function) by cross-correlation. This, in turn, also aligned the Sla1-GFP and Rvs167-GFP trajectories. Once all the raw trajectories were aligned, we computed an average binning of all the time points within 0.5 s intervals (‘average_dw’ function). Figure 5D shows the average for the fimbrin-mCherry trajectories imaged together with Sla1-GFP. The curves would be identical for the fimbrin-mCherry trajectories imaged with Rvs167-GFP.

All lifetime data are stored on https://github.com/apicco/EvoCellBio under Protein_lifetimes and the Python3 (https://www.python.org/downloads/) scripts used to generate the plots.

Spot dynamics were tracked with ParticleTracker (v1.5, Sblazarini and Koumoutsakos, 2005) and analysed with the trajalin python module distribution (https://apicco.github.io/trajectory_alignment/, Picco and Kaksonen, 2017 Methods in Cell Biology) and Python3. All the trajectories and the scripts used to generate the plots are stored on https://github.com/apicco/EvoCellBio under Protein_tracking.

The WASP fluorescence intensities were determined using PaticleTracker and aligned by cross-correlating the fluorescence intensity profiles of fimbrin-mCherry. All the data and the Python3 scripts used to generate the plots in Figure 7 are available on https://github.com/apicco/EvoCellBio under Protein_FI/WASP.

## SUPPLEMENTAL FIGURES

**Supplemental Figure S1:**
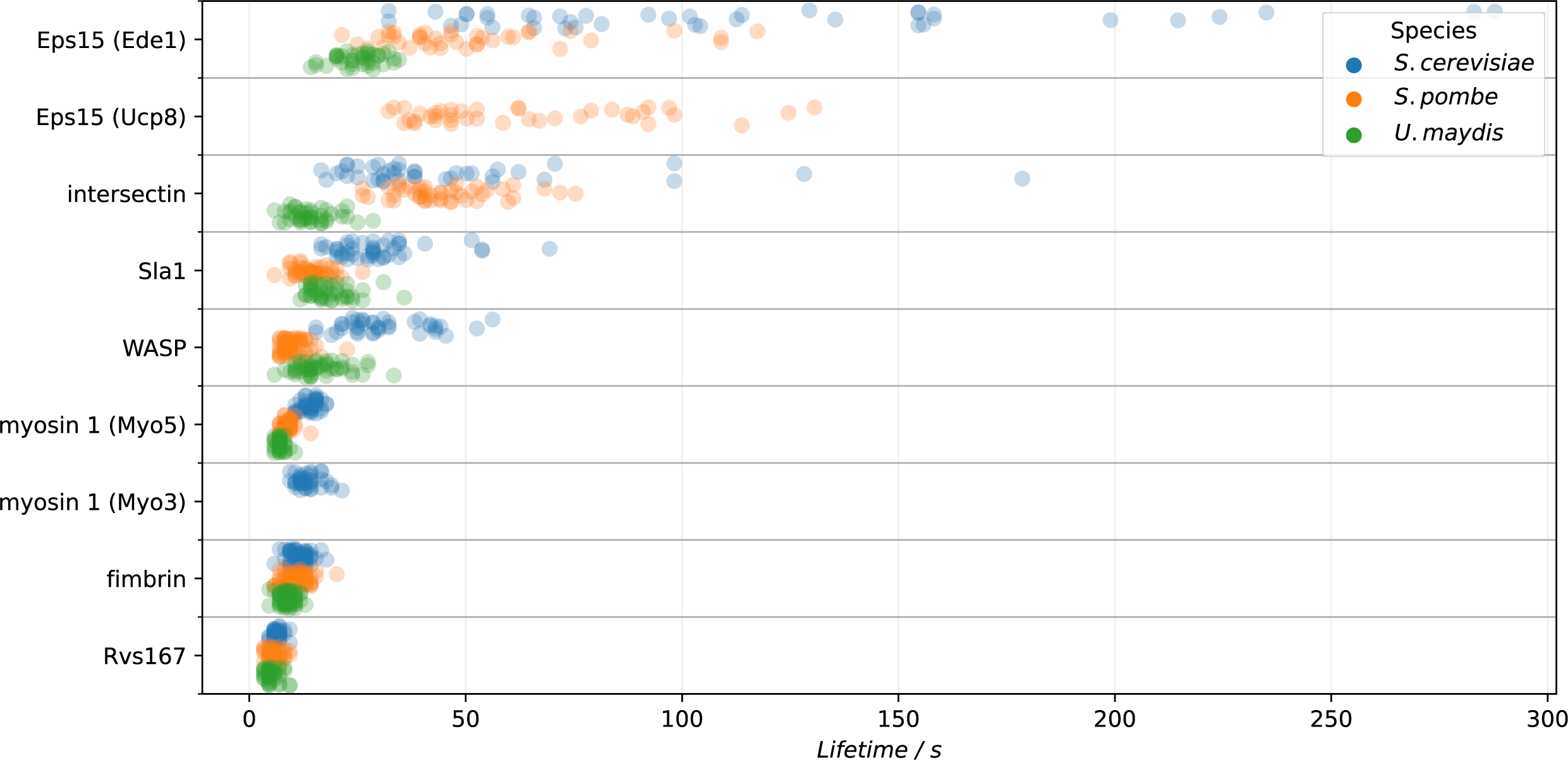
The individual lifetimes measured for each protein in each species to generate Figure 4.

**Supplemental Figure S2:**
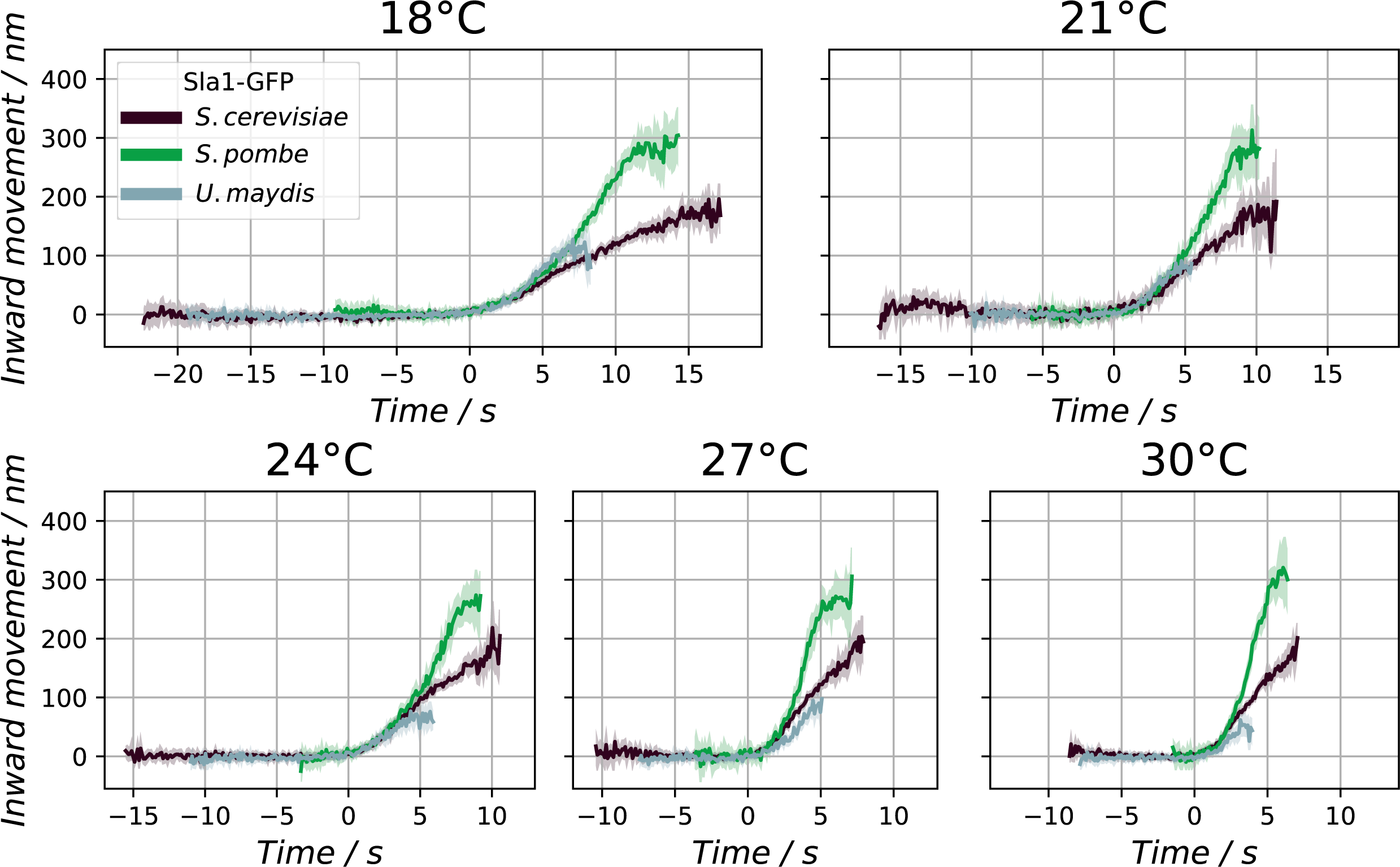
The complete average trajectories detailing the invagination dynamics of the coat marker Sla1-GFP in *S. cerevisiae*, *S. pombe*, and *U. maydis* at 18°C, 21°C, 24°C, 27°C, and 30°C.

## SUPPLEMENTAL TABLES

**Supplemental Table S1:** Endocytic protein lifetimes.

|  | Protein | Lifetime / s | SD / s |
| --- | --- | --- | --- |
| *S. cerevisiae* | Ede1 | 111.21 | 66.48 |
| *S. cerevisiae* | Pan1 | 44.72 | 31.06 |
| *S. cerevisiae* | Sla1 | 28.69 | 9.44 |
| *S. cerevisiae* | Wasp | 30.12 | 9.41 |
| *S. cerevisiae* | Myo3 | 12.48 | 2.67 |
| *S. cerevisiae* | Myo5 | 12.96 | 1.92 |
| *S. cerevisiae* | Rvs167 | 5.41 | 1.76 |
| *S. cerevisiae* | Fim1 | 10.54 | 2.2 |
| *S. pombe* | Ucp8 | 62.4 | 27.6 |
| *S. pombe* | Ede1 | 50.94 | 23.65 |
| *S. pombe* | Pan1 | 43.96 | 11.3 |
| *S. pombe* | Sla1 | 13.41 | 3.35 |
| *S. pombe* | Wasp | 9.09 | 2.8 |
| *S. pombe* | Myo1 | 7.8 | 1.66 |
| *S. pombe* | Rvs167 | 4.76 | 2.21 |
| *S. pombe* | Fim1 | 9.69 | 2.89 |
| *U. maydis* | Ede1 | 24.15 | 5.96 |
| *U. maydis* | Pan1 | 13.37 | 4.77 |
| *U. maydis* | Sla1 | 17.69 | 4.87 |
| *U. maydis* | Wasp | 15.4 | 5.45 |
| *U. maydis* | Myo1 | 6.12 | 0.96 |
| *U. maydis* | Rvs167 | 4.19 | 2.54 |
| *U. maydis* | Fim1 | 7.82 | 1.63 |

Endocytic protein lifetime values obtained from TIRF movies.

**Supplemental Table S2:** Media formulations.

| Media | Species | Components (w/v) |
| --- | --- | --- |
| YPD | *S.c.* | 1% Yeast extract, 2% Bacto peptone, 2% Glucose |
| YES | *S.p.* | 0.5% Yeast extract, 3% Glucose, 0.0225% Adenine, 0.0225% Histidine, 0.0225% Leucine, 0.0225% Lysine, 0.0225% Uracil |
| YEPS-light | *U.m.* | 1% Yeast extract, 0.4% Bacto peptone, 0.4% Sucrose |
| SC-Trp | *S.c.* | 0.67% Yeast nitrogen base w/o Folic acid w/o Riboflavin, 2% Glucose. 0.192% SC-Trp |
| EMM | *S.c.*<br>*S.p.*<br>*U.m.* | 0.3% Potassium phthalate, monobasic, 0.22% Na<sub>2</sub>HPO<sub>4</sub>*2H<sub>2</sub>O, 0.5% NH<sub>4</sub>Cl, 2% Glucose, 1X 'Salts solution', 1X 'Vitamins solution', 1X 'Minerals solution'<br><u>50 X Salts solution:</u> 5.28% MgCl<sub>2</sub>*6H<sub>2</sub>O, 0.0734% CaCl<sub>2</sub>*2H<sub>2</sub>O, 5% KCl, 0.2% Na<sub>2</sub>SO<sub>4</sub><br><u>1,000X Vitamins solution:</u> 0.1% D-Pantothenate, 1% Nicotinic acid, 1% myo-Inositol, 0.001% Biotin<br><u>10,000X Minerals solution:</u> 0.5% Boric acid, 0.4% MnSO<sub>4</sub>, 0.4% ZnSO<sub>4</sub>*7H<sub>2</sub>O, 0.2% FeCl<sub>2</sub>*6H<sub>2</sub>O, 0.04% MoO<sub>3</sub>, 0.1% KI, 0.04% CuSO<sub>4</sub>*5H<sub>2</sub>O, 1% Citric acid |

Media used in this study. For plates 2% agar was added to these media.

**Supplemental Table S3:** Plasmid table.

| Plasmid | Source | Description |
| --- | --- | --- |
| pMK0003 | M. Kaksonen | C-terminal eGFP::HIS3 for *S. cerevisiae* |
| pMK0005 | M. Kaksonen | C-terminal mCherry:KanMX for *S. cerevisiae* |
| pSM1685 | Sophie Martin | C-terminal sfGFP for *S. pombe* |
| pSM0684 | Sophie Martin | C-terminal mCherry for *S. pombe* |
| pUMa1694<br>pStorl_5-1n<br>cTerm-Gfp_natR | Michael<br>Feldbrügge | BsaI Gateway Storage plasmid C-terminal eGFP for *U. maydis* |
| pUMA1557<br>pMK0384 | This study | BsaI Gateway Storage plasmid C-terminal mCherry for *U. maydis* |
| pUMa2691<br>storage<br>Sapl cTerm-Gfp_natR | Michael<br>Feldbrügge | SapI Gateway Storage plasmid C-terminal eGFP or *U. maydis* |
| pUMa1467<br>pDest BsaI | Michael<br>Feldbrügge | BsaI Gateway Destination plasmid ClonNAT resistance for *U. maydis* |
| pUMa2074_<br>pUC57<br>AmpR SapI | Michael<br>Feldbrügge | SapI Gateway Destination plasmid Hygromycin resistance or *U. maydis* |

**Supplemental Table S4:** Strain table.

| Name | Species | Genotype | Source |
| --- | --- | --- | --- |
| MKY0104 | *S.c.* | *MATa/MATα, his3-Δ200/his3-Δ200, leu2-3,112/leu2-3,112, ura3-52/ura3-52, lys2-801/+* | Kaksonen lab |
| MKY2833 | *S.c.* | *MATa, his3-Δ200, leu2-3,112, ura3-52, lys2-801, SLA1-eGFP::HIS3MX6* | Kaksonen lab |
| MKY3863 | *S.c.* | *MATα, his3-Δ200, leu2-3,112, ura3-52, lys2-801, SAC6-eGFP::HIS3MX6* | Kaksonen lab |
| MKY3996 | *S.c.* | *MATα, his3-Δ200, leu2-3,112, ura3-52, lys2-801, SAC6-mCherry::kanMX4* | Kaksonen lab |
| MKY4014 | *S.c.* | *MATα, his3-Δ200, leu2-3,112, ura3-52, lys2-801, SLA1-eGFP::HIS3MX6, SAC6-mCherry::kanMX4* | Kaksonen lab |
| MKY4015 | *S.c.* | *MATa, his3-Δ200, leu2-3,112, ura3-52, lys2-801, SLA1-eGFP::HIS3MX6, SAC6-mCherry::kanMX4* | Kaksonen lab |
| MKY4017 | *S.c.* | *MATa, his3-Δ200, leu2-3,112, ura3-52, lys2-801, PAN1-eGFP::HIS3MX6, SAC6-mCherry::kanMX4* | Kaksonen lab |
| MKY4019 | *S.c.* | *MATa, his3-Δ200, leu2-3,112, ura3-52, lys2-801, RVS167-eGFP::HIS3MX6, SAC6-mCherry::kanMX4* | Kaksonen lab |
| MKY4021 | *S.c.* | *MATa, his3-Δ200, leu2-3,112, ura3-52, lys2-801, LAS17-eGFP::HIS3MX6, SAC6-mCherry::kanMX4* | Kaksonen lab |
| MKY4155 | *S.c.* | *MATa, his3-Δ200, leu2-3,112, ura3-52, lys2-801, ARC18-myeGFP::NAT, SAC6-mCherry::kanMX4* | Kaksonen lab |
| MKY4390 | *S.c.* | *MATα, his3-Δ200, leu2-3,112, ura3-52, lys2-801, las17Δ::SpWSP1-eGFP::HIS3MX6, SAC6-mCherry::kanMX4* | Kaksonen lab |
| MKY4612 | *S.c.* | *MATa, his3-Δ200, leu2-3,112, ura3-52, lys2-801, EDE1-eGFP::NAT, SAC6-mCherry::kanMX4* | Kaksonen lab |
| MKY4649 | *S.c.* | *MATa, his3-Δ200, leu2-3,112, ura3-52, lys2-801, MYO5-eGFP::HIS3MX6, SAC6-mCherry::kanMX4* | Kaksonen lab |
| MKY4651 | *S.c.* | *MATa, his3-Δ200, leu2-3,112, ura3-52, lys2-801, MYO3-eGFP::HIS3MX6, SAC6-mCherry::kanMX4* | Kaksonen lab |
| MKY4794 | *S.c.* | *MATα, his3-Δ200, leu2-3,112, ura3-52, lys2-801, sla1Δ::URA3, las17Δ::SpWSP1-eGFP::HISMX6, SAC6-mCherry::kanMX4* | Kaksonen lab |
| MKY4801 | *S.c.* | *MATα, his3-Δ200, leu2-3,112, ura3-52, lys2-801, sla1Δ::URA3, LAS17-eGFP::HISMX6, SAC6-mCherry::kanMX4* | Kaksonen lab |
| MKYP0003 (YSM1371) | *S.p.* | *h+* | Sophie Martin |
| MKYP0020 | *S.p.* | *h-* | Kaksonen lab |
| MKYP0006 | *S.c.* | *h+, SHD1-sfGFP::kanMX6* | Kaksonen lab |
| MKYP0010 | *S.c.* | *h+, FIM1-sfGFP::kanMX6* | Kaksonen lab |
| MKYP0021 | *S.c.* | *h-, FIM1-mCherry::natMX6* | Kaksonen lab |
| MKYP0022 | *S.c.* | *h-, SHD1-sfGFP::kanMX6, FIM1-mCherry::natMX6* | Kaksonen lab |
| MKYP0023 | *S.p.* | *h+, HOB1-sfGFP::kanMX6, FIM1-mCherry::natMX6* | Kaksonen lab |
| MKYP0024 | *S.p.* | *h+, PAN1-sfGFP::kanMX6, FIM1-mCherry::natMX6* | Kaksonen lab |
| MKYP0026 | *S.p.* | *h-, WSP1-sfGFP::kanMX6, FIM1-mCherry::natMX6* | Kaksonen lab |
| MKYP0039 | *S.p.* | *h+, MYO1-sfGFP::kanMX6, FIM1-mCherry::natMX6* | Kaksonen lab |
| MKYP0068 | *S.p.* | *h-, ARC3-5(GA)-sfGFP::kanMX6, FIM1-mCherry::natMX6* | Kaksonen lab |
| MKYP0091 | *S.p.* | *h-, EDE1-sfGFP::kanMX6, FIM1-mCherry::natMX6* | Kaksonen lab |
| MKYP0092 | *S.p.* | *h-, UCP8-sfGFP::kanMX6, FIM1-mCherry::natMX6* | Kaksonen lab |
| MKUM0000 | *U.m.* | *a1 b1* | Fungal |
| (UM521) |  |  | Genetic Stock Center |
| MKUM0001 | *U.m.* | *a1 b1 SLA1-eGFP::Nat<sup>R</sup>* | Kaksonen lab |
| MKUM0005 | *U.m.* | *a1 b1 FIM1-eGFP::Nat<sup>R</sup>* | Kaksonen lab |
| MKUM0006 | *U.m.* | *FIM1-mCherry::Hph<sup>R</sup>* | Kaksonen lab |
| MKUM0015 | *U.m.* | *a1 b1 EDE1-eGFP::Nat<sup>R</sup> FIM1-mCherry::Hph<sup>R</sup>* | Kaksonen lab |
| MKUM0020 | *U.m.* | *a1 b1 PAN1-eGFP::Nat<sup>R</sup> FIM1-mCherry::Hph<sup>R</sup>* | Kaksonen lab |
| MKUM0014 | *U.m.* | *a1 b1 SLA1-eGFP::Nat<sup>R</sup> FIM1-mCherry::Hph<sup>R</sup>* | Kaksonen lab |
| MKUM0024 | *U.m.* | *a1 b1 WSP1-eGFP::Nat<sup>R</sup> FIM1-mCherry::Hph<sup>R</sup>* | Kaksonen lab |
| MKUM0016 | *U.m.* | *a1 b1 MYO1-eGFP::Nat<sup>R</sup> FIM1-mCherry::Hph<sup>R</sup>* | Kaksonen lab |
| MKUM0019 | *U.m.* | *a1 b1 RVS167-eGFP::Nat<sup>R</sup> FIM1-mCherry::Hph<sup>R</sup>* | Kaksonen lab |
| MKUM0011 | *U.m.* | *a1 b1 ARC3-eGFP::Nat<sup>R</sup> FIM1-mCherry::Hph<sup>R</sup>* | Kaksonen lab |

Strains used in this study

## ACKNOWLEDGEMENTS

We thank Sophie Martin and Vincent Vincenzetti for *S. pombe* strains and advice and Michael Feldbrügge and Karl Haag for *U. maydis* strains and advice. We thank Markku Hakala, Anne-Laure Boinet, Mamta, Yushi Jiang, Nastasia Verdes, and Bagyashree V. T. for critical reading of the manuscript. This work was supported by the Swiss National Science Foundation grant (310030_212288).

